# ViruDB: An Assay-Aware Molecular Database and Endpoint-Adaptive Multimodal Learning Framework for Antibiofilm, Quorum-Sensing Inhibition, and Antivirulence Research

**DOI:** 10.64898/2026.11.21.768264

**Authors:** Pankaj Paul, Ruchita Santra

## Abstract

Antibiofilm and quorum-sensing inhibitory molecules have become increasingly attractive candidates to complement traditional antibacterial approaches, but the experimental evidence ranges from heterogeneous assays, organisms, methods of activity measurement, and chemical representation. We created ViruDB, a molecular evidence resource that is assay-aware, and tested an endpoint-adaptive multimodal learning framework for quantitative activity prediction. Curated collection of 8765 experimental evidence records for 3563 unique molecules: 2,959 quantitative observations, of which 2,345 were antibiofilm, 569 were quorum-sensing inhibition [QSI], and 45 were antivirulence. Molecular representations included 217 2D descriptors from RDKit, 2,048-bit ECFP4 fingerprints, molecular graphs, and some information about the biological context. A molecule- and scaffold-disjoint primary partition was used, which resulted in 2,339 observations in the training set, 265 observations in the validation set and 355 observations in the independent test set. ViruFusion-TaskExpert-MTL and an endpoint specific combination with classical regressors were selected by validation. The final predictor yielded an RMSE value of 0.5460 and an R^2^ of 0.4816 on the antibiofilm activity dataset (*n*=286) and an RMSE value of 0.5744 and an R^2^ of 0.4082 on the QSI dataset (*n*=67) in the reported frozen independent evaluation. Exploratory – anti-virulence regression (*n*=2 test observations). The results validate quantitative, endpoint based prioritization under chemical novelty and highlight the significance of assay provenance, assay leakage control and external biological assay validation.

## 1. INTRODUCTION

### 1.1. Antimicrobial resistance and biofilm-associated infection

Due to the growing spread of antimicrobial resistance, there has been renewed interest in strategies that are not solely dependent on killing or slowing the growth of bacteria [1–3]. Biofilms promote persistent infections by creating external matrices, changing metabolic functions, creating spatial heterogeneity, and increasing resistance to antimicrobials. It is important to note that the two types of tolerance and antimicrobial resistance that are associated with biofilms can be separated, though they can occur at the same time. If a computational programme is to be developed to search for antibiofilm molecules, then phenotypic definitions that differentiate prevention of biofilm from disruption of a mature biofilm and ordinary growth inhibition are required [4–8].

Antivirulence interventions aim to disable pathogenic aspects (adhesion, quorum sensing, toxin release, motility, secretion), but do not necessarily kill the organism. They possess the potential to change evolutionary selection but are context dependent – resistance to antivirulence agents is possible [1,2]. In turn, predictions of anti-biofilm or quorum-sensing activity were not necessarily read as an experimental anchor for molecular phenotypes, but rather as evidence of the therapeutic effectiveness [9–11].

### 1.2. Quorum-sensing pathways and anti-virulence targets

The ability to communicate, or “talk to each other,” is known as quorum sensing (QS) and links population dependent signals to transcriptional programmes. Acyl-homoserine lactone systems such as the *Las* and *Rhl* circuits of *P. aeruginosa* control a variety of secreted factors and collective behaviours in Gram-negative bacteria. In addition, there are other mechanisms, including the *Pseudomonas* quinolone signal system, communication with AI-2, and Gram-positive peptide-based communication systems like *Agr* [12,13]. Phenotypes associated with QS interference are reported activity, pyocyanin production, elastase, protease, rhamnolipid expression, motility and biofilm formation. In the ViruDB evidence framework (Fig. 1), however, the difference between the phenotypic inhibition of QS and the direct binding to *LasR*, *RhlR*, or *PqsR* is maintained [14–18].

**Fig. 1.**
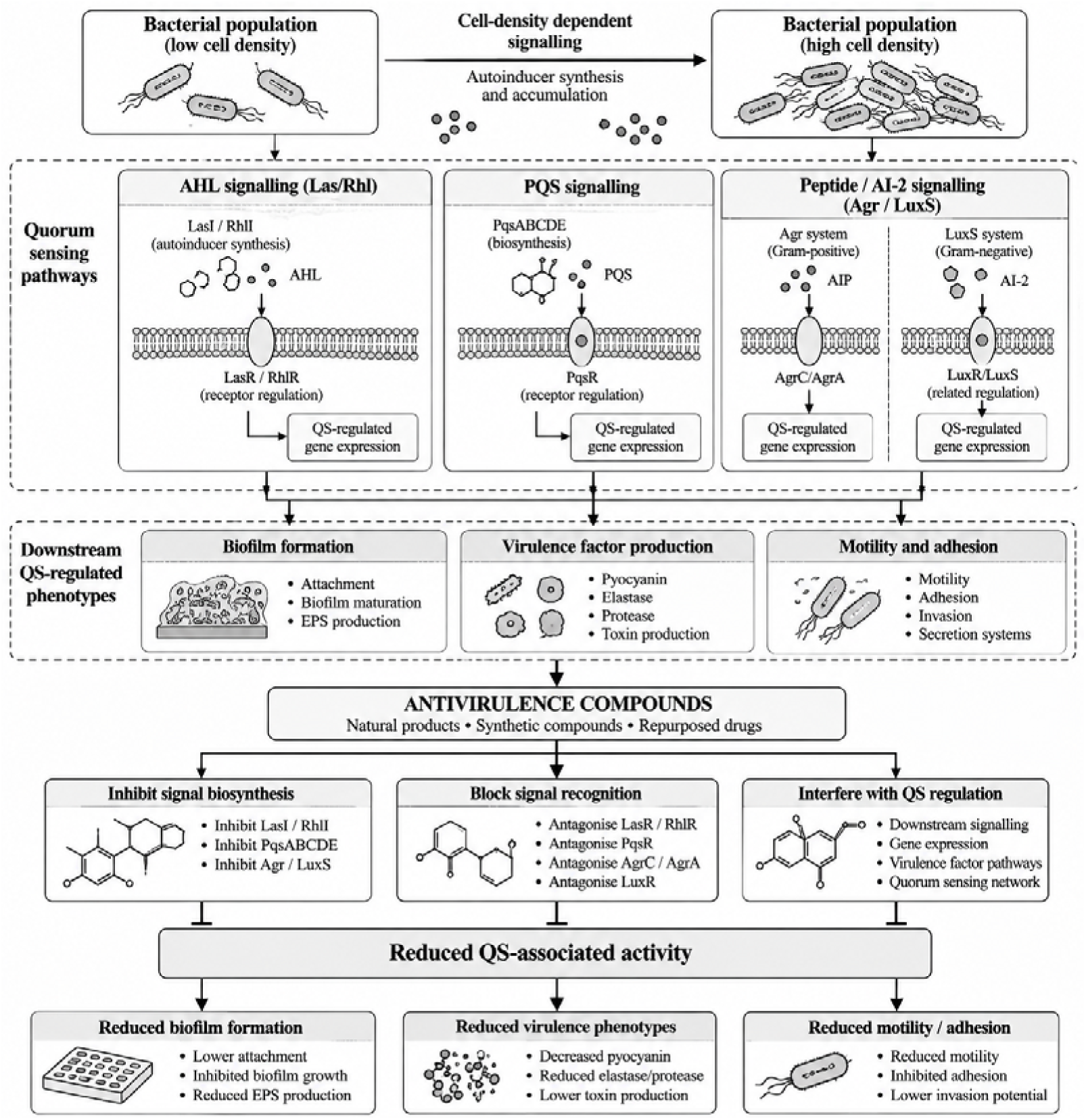
Biological mechanisms of quorum-sensing regulation and potential anti-virulence intervention.

### 1.3. Chemical diversity and the need for assay-aware resources

There is a wide range of literature on phenolics, flavonoids, coumarins, alkaloids, terpenoids, microbial metabolites, marine natural products, and synthetic analogues all studied in the field of anti-biofilm and QS. However, membership in a chemical class doesn’t prove inhibitory activity. Results will vary depending on bacterial strain, medium, concentration, time of exposure, readout of the assay, and controlled growth of the bacteria. The quantitative structure–activity relationship (QSAR) model, therefore, necessitates reliable molecular identity as well as a defensible mapping between experimental measurement and numerical target variable [15,18].

### 1.4. Those that are available, and what remains to be done

aBiofilm offers antibiofilm-agent information and prediction-oriented functionality [19]; Molib and Biofilm-i offer prediction-oriented functionality for antibiofilm-related outcomes based on machine learning [20] and [21]; Anti-Biofilm offers quantitative IC_50_-oriented modelling [22]. While ChEMBL and PubChem offer wide-ranging chemical and assay infrastructure [23,24], Quorumpeps concentrates on signalling peptides [25]. The 2025 version of VFDB also includes 902 antiseptic compounds, which are derived from 262 studies [26] and are used to prevent infection from developing. So, ViruDB is not a first antivirulence compound database, but rather a multi-endpoint platform that offers assay-informed quantitative evidence and leakage-controlled predictive evaluation. Table 1 places ViruDB in this context.

**TABLE 1.** Positioning of ViruDB among representative resources.

| **Resource** | **Primary focus** | **Prediction scope** | **Distinctive limitation / boundary** |
|---|---|---|---|
| aBiofilm [19] | Antibiofilm agents | Antibiofilm classification/tools | Not a unified quantitative three-endpoint benchmark |
| Molib [20] | Biofilm inhibitory molecules | ML classification | Biofilm-focused |
| Biofilm-i [21] | Biofilm inhibitor QSAR | Regression and web platform | Biofilm-focused endpoints |
| Anti-Biofilm [22] | Biofilm IC<sub>50</sub> | ML regression | IC<sub>50</sub>-focused biofilm prediction |
| Quorumpeps [25] | QS peptides | Peptide information | Not general small-molecule QSI |
| VFDB 2025 [26] | Virulence factors and compounds | Knowledge resource | Not the ViruDB frozen multi-endpoint predictor |
| ChEMBL [23] | Broad bioactivity | Data resource | Heterogeneous assay labels |
| PubChem [24] | Chemical and bioassay records | Data resource | Broad non-specific assays |
| ViruDB (this work) | Assay-aware ABF/QSI/AV | Endpoint-adaptive regression | AV prediction remains exploratory |

The unique value added is not the first collection of Antivirulence compounds, but the ability to integrate with evidence provenance at the molecule level, quantitative eligibility by assay and compound endpoint specific evaluation. While there are several broad bioactivity repositories that provide chemical information and experimental data, the ViruDB benchmark explicitly distinguishes data from antibiofilm, QSI and antivirulence effects, preserves the context of the assays, and applies a prediction policy selected for validation and tests it in the molecule/scaffold separation mode. The role of the database contribution is not alternative to the prediction contribution, but supplementary, helping to retrieve evidence and to provide evidence of the traceability; the role of the prediction contribution is to assess a defined quantitative modelling subset.

### 1.5. Research objectives and contributions

This study aimed to answer four related questions: (i) if it is possible to standardize anti-virulence observations while avoiding merging biologically different assays; (ii) how molecule/scaffold-disjoint performance of molecular descriptors, fingerprints, and graph representations; (iii) if task-specific experts can help decrease negative transfer between antibiofilm and QSI regression; and (iv) if validation-selected endpoint adaptation can help improve independent-test prediction without tuning on test labels. They include a molecular portfolio with evidence-based molecules, a quantitative three endpoint benchmark, a multimodal task-expert learning architecture, and a frozen endpoint-specific prediction policy. The study concludes that a publicly deployed interface, ADMET or docking was experimentally completed.

## 2. MATERIALS AND METHODS

### 2.1. Overall experimental design

This study involved chemical and biological evidence curation, quantitative eligibility evaluation, molecular feature engineering, leakage-controlled data partitioning, conventional QSAR benchmarks, comparison of neural architectures, selection of endpoint-specific policies and one frozen independent evaluation. The entire process is summarized in Fig. 2. There was one model development output investigated, the final frozen metrics were from the full final optimization execution log, not the diagnostic workbook.

**Fig. 2.**
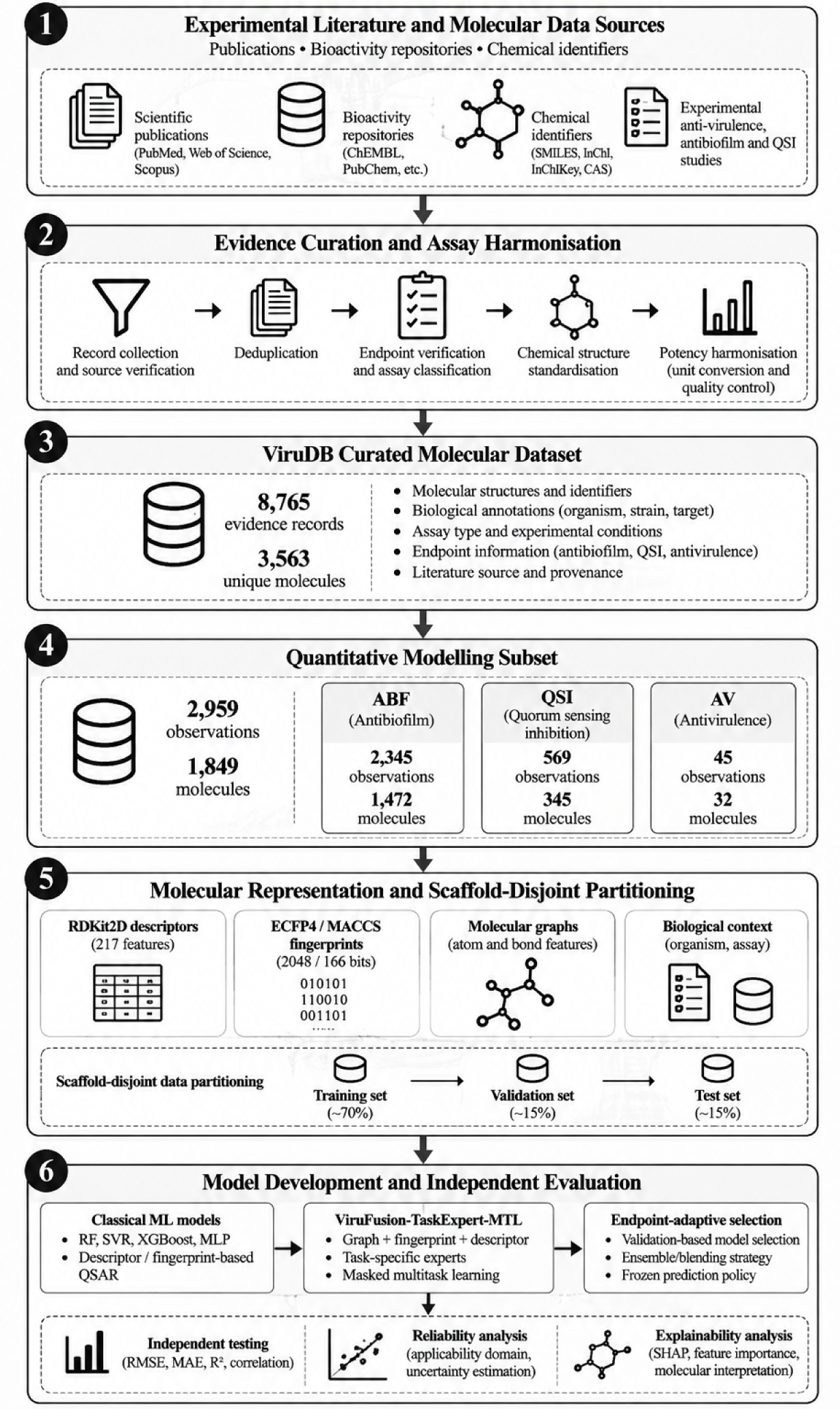
End-to-end workflow linking source evidence, molecular curation, quantitative modelling, and independent evaluation.

### 2.2. Evidence acquisition, molecular identity, and endpoint harmonization

The curated master file contains molecule identifiers, canonical SMILES and InChIKeys, physicochemical properties, Bemis–Murcko scaffolds, pathogen annotations, assay organism, mechanism grouping, measurement type and units, transformed activity (if applicable), bibliographic identifiers, quality indicators and temporal or model-eligibility flags. Multiple observations of the same compound were considered as separate molecules, if they were assays and/or biological conditions that were different.

Provenance and assay data for source and assay was represented by bibliographic identifiers, molecular identifiers, organism annotation, measurement definition, reported units and quantitative eligibility flags in ViruDB. The curated subset, which included all the evidence records, consisted of 8,765 evidence records, with 2,959 evidence records being part of the quantitative modelling subset, and 5,806 evidence records being retained as non-model-ready evidence records (Table 2). These numbers reflect the output of the completed curation; no quantification of source-specific retrieval and previous exclusion numbers are available in the overall dataset summary and are therefore not cited.

**TABLE 2.**
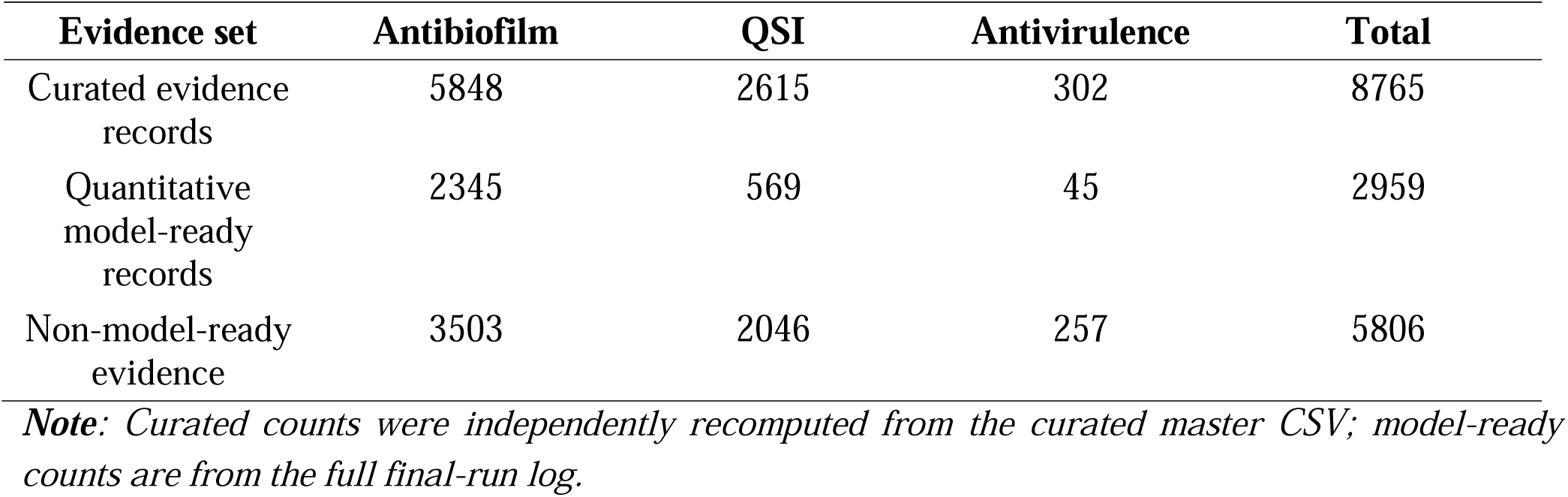
Observed curation totals and endpoint coverage.

| **Evidence set** | **Antibiofilm** | **QSI** | **Antivirulence** | **Total** |
|---|---|---|---|---|
| Curated evidence records | 5848 | 2615 | 302 | 8765 |
| Quantitative model-ready records | 2345 | 569 | 45 | 2959 |
| Non-model-ready evidence | 3503 | 2046 | 257 | 5806 |

***Note****: Curated counts were independently recomputed from the curated master CSV; model-ready counts are from the full final-run log.*

Activity harmonization considered each measurement in its own assay context, instead of viewing it as interchangeable potency label. IC_50_ is the concentration that inhibits 50% of the defined assay; MIC = concentration that inhibits visible growth; MBIC = concentration that inhibits the growth of an established biofilm under the defined assay; and MBEC = concentration that eliminates an established biofilm under the defined assay. Single dose percent inhibition not transformed to IC_50_ due to lack of appropriate concentration-response. The molar concentrations of the eligible compounds were expressed as pActivity = −log_10_(C/mol.L^−1^); a mass concentration was converted to a molar concentration only if a clear molecular mass and unit of reporting were identified. The original ‘readout’, ‘assay type’, ‘organism’ and ‘units’ were retained.

The original measurement labels were retained and the only transformations on concentrations were of chemical meaning, such as molar numbers. Potency was indicated as pActivity, which is equivalent to −log_10_(C in mol.L^−1^) when appropriate. The terms percentage inhibition, MIC, MBIC, MBEC, and mechanistically different readout were not taken for granted to be interchangeable. The present curated dataset is not a confirmation of molecular target engagement for each compound. Algorithm 1 summarizes the evidence-curation procedure.

**Algorithm 1.** Evidence-to-quantitative record curation

1. Load experimental records with source and assay metadata.
2. Resolve molecule identity; canonicalize structure and compute identifiers.
3. Classify phenotype and measurement type; preserve original units.
4. Reject unresolved, incompatible, or duplicated evidence according to logged rules.
5. Transform only eligible molar potency measurements; retain provenance.
6. Assign curated evidence and model-eligibility flags; audit outputs.

### 2.3. Molecular representations and data composition

There are 8,765 evidence records in the curated collection and 3,563 unique molecular structures. The final quantitative subset consists of 2,959 observations for 1,849 individual molecules. The distribution is highly skewed, with just 45 quantitative observations carried out for the antivirulence task as summarized in Table 3. This imbalance resulted in task weighting and conservative reporting.

**TABLE 3.** Quantitative dataset and endpoint imbalance.

| **Endpoint** | **Model-ready *n*** | **Share of 2,959** | **Interpretation** |
|---|---|---|---|
| Antibiofilm | 2345 | 79.25% | Primary regression endpoint |
| QSI | 569 | 19.23% | Primary regression endpoint |
| Antivirulence | 45 | 1.52% | Exploratory auxiliary endpoint |
| Total | 2959 | 100% | 1,849 distinct molecules |

There are two sources of information: structure-related feature branches and biological-context. The descriptors derived from RDKit, ECFP4, MACCS keys, and molecular graph are based on the standardized molecular structure, while the organism identifiers and the endpoint identifiers are based on curated experimental metadata. This distinction is important because it disallows biological assay labels from being incorrectly said to be intrinsic molecular characteristics. Results of the quantitative subset are reported separately from task-level sample counts and test coverage, even though most of the observations are of the antibiofilm type (79.25%).

A total of 217 2D descriptors from RDKit, 2,048-bit ECFP4/Morgan fingerprints (radius 2), 166 classical features in the feature audit, and molecular graphs generated from sanitized structures were used for featurization. The evaluation graph cache was frozen for 1,849 molecules with a dimensionality of 28 atoms and 35 organism identifiers and seven endpoint identifiers in the model context vocabulary. The representation families are shown in Table 4. These features convey the context of the assay as well as encoding molecular structure but are not indicative of experimental evidence of target binding.

**TABLE 4.** Molecular and contextual representation families.

| **Representation** | **Dimension** | **Source** | **Role** |
|---|---|---|---|
| RDKit 2D | 217 | Canonical molecular structures | Physicochemical topology |
| ECFP4 | 2,048 bits | Morgan radius 2 | Substructure fingerprint |
| MACCS | 166 bits | Molecular keys | Classical benchmark |
| Molecular graph | Variable; atom features=28 | Atoms and bonds | Graph message passing |
| Organism context | 35 categories | Curated organism vocabulary | Biological context |
| Endpoint context | 7 categories | Curated endpoint vocabulary | Assay context |

### 2.4. Chemical-space and scaffold characterization

The measured properties (RDKit) as well as the distribution of organisms and frequency of non-empty Bemis–Murcko scaffolds within the curated master were used in chemical space analysis. These analyses do not depict the distribution of independent-test predictions. The quantitative modelling subset was kept separate when computing task-level metrics. The chemical-space audit revealed representation imbalance, scaffold families, and the level of scaffold-level separation to investigate generalization [27].

### 2.5. Molecule- and scaffold-disjoint splitting

The final Primary_Multitask_Split split the data into 2,339 training observations, 265 validation observations, and 355 test observations. There were no conflicts found in all three pairwise comparisons between Train, Validation and Test for both molecule identifiers and scaffolds. Table 5 provides end point counts. These limitations decrease the likelihood of evaluating the model on replicates or highly similar families of scaffold structures that were used for training, but do not completely rule out any type of assay, source, or target leakage. This split was corrected prior to the selection of neural model and blend (Algorithm 2) [28].

**TABLE 5.** Frozen primary partition and molecule/scaffold integrity.

| **Endpoint** | **Train** | **Validation** | **Test** | **Total** |
|---|---|---|---|---|
| Antibiofilm | 1871 | 188 | 286 | 2345 |
| QSI | 444 | 58 | 67 | 569 |
| Antivirulence | 24 | 19 | 2 | 45 |
| Total | 2339 | 265 | 355 | 2959 |
| Molecule/scaffold overlaps | 0 | 0 | 0 | Pairwise zero |

All records with the same canonical molecule or scaffold were given leakage control. Virtually identical observations of a compound (from different sources or endpoints) were placed in the same partition, and the scaffold structures were likewise contained within partition boundaries. This audit identified no pairwise overlaps in molecule and scaffold identifiers (Table 5). This structural integrity check decreased the likelihood of duplicate-structure leakage, but there was no independent evaluation for source-level or assay-protocol or temporal dependences. This audit is based on molecule keys, scaffold definitions, split assignments and allocation rules.

**Algorithm 2.** Primary scaffold-disjoint split and audit

1. Assign each quantitative record a canonical molecule key and Murcko scaffold.
2. Group molecules and scaffolds so groups cannot cross partitions.
3. Allocate training, validation, and independent-test records under endpoint constraints.
4. Verify all pairwise intersections of molecule and scaffold identifiers are empty.
5. Fit preprocessing and estimators using training records only.
6. Preserve validation for model/policy selection and lock independent test.

### 2.6. Classical QSAR benchmarks

Dummy, Ridge, k-nearest neighbours, support vector regression (SVR), Random Forest (RF), Extra Trees, XGBoost and multilayer perceptron were the conventional regressors used. Candidate feature representations included RDKit descriptors, ECFP4, MACCS, and a few hybrids of these. The training and validation were split from the independent test. The classical models chosen for final same-split reference were ECFP4/RF for antibiofilm, ECFP4/SVR for QSI, and ECFP4/RF for antivirulence. The latter was kept as a comparison on an exploratory basis. The full tuning grid is not reported in this document [29–31] and is omitted from Table 6 which specifies the scope for the benchmark.

**TABLE 6.** Classical benchmark and selection configuration.

| **Model family** | **Examples** | **Feature groups** | **Selection basis** |
|---|---|---|---|
| Linear / simple | Dummy, Ridge, KNN | RDKit2D, ECFP4, MACCS | Training/CV and validation |
| Kernel-based | SVR | ECFP4 and alternatives | Validation RMSE |
| Ensembles | RF, ExtraTrees, XGBoost | Descriptors/fingerprints/hybrid | Validation RMSE |
| Neural baseline | MLP | Descriptor/fingerprint features | Validation RMSE |
| Final classical comparator | RF (ABF), SVR (QSI), RF (AV) | ECFP4 | Same primary split |

Additional complexity of multimodal learning did not result in an increase of prediction error, as established by classical benchmarks. Simple ridge and dummy regressors were used as baselines, kernel-based nonlinear SVR as a baseline, and RF, Extra Trees, and XGBoost as baseline ensemble methods. The same primary partition, endpoint-specific response labels and test metric definitions were used in the reported final comparisons. Table 7 shows same-split comparisons for antibiofilm and QSI capabilities available; no performance claims have been made outside the candidate grid without the results for a particular configuration.

**TABLE 7.** Verified same-split classical reference models.

| **Endpoint** | **Classical comparator** | **Representation** | **Test n** | **Test RMSE** |
|---|---|---|---|---|
| Antibiofilm | Random Forest | ECFP4 | 286 | 0.6373 |
| QSI | SVR | ECFP4 | 67 | 0.5794 |
| Antivirulence | Random Forest | ECFP4 | 2 | Not reported in the frozen evaluation summary |

***Note:*** *The classical reference RMSEs were obtained from the reported same-split evaluation. Additional baseline metrics are not tabulated because their complete numerical outputs were not included in the evaluated summary.*

**TABLE 8.** Validation-only neural architecture and frozen endpoint policy.

| **Architecture / endpoint** | **Validation RMSE** | **Selection / neural weight** |
|---|---|---|
| TaskExpert-MTL primary score | 0.6450 | Selected architecture |
| GatedMoE-MTL primary score | 0.6605 | Not selected |
| Antibiofilm final blend | 0.5583 | Neural 0.50; RF 0.50 |
| QSI final policy | 0.6529 | Neural 1.00 |
| Antivirulence exploratory blend | 0.7134 | Neural 0.30; RF 0.70 |

### 2.7. ViruFusion-TaskExpert-MTL architecture

ViruFusion was developed to merge graph-derived representations, fingerprint, descriptor and biological-context embeddings. Connectivity is captured in molecular graph information, local substructure patterns and descriptors provide global physicochemical summaries, and circular fingerprints capture local patterns. The fusion representation goes through the task-specific experts, and the endpoint regression heads (Fig. 3). A masked loss assigns a loss to only those tasks for which labels are available. Task-loss weights were 1.00 for antibiofilm, 1.25 for QSI and 0.35 for antivirulence in the frozen evaluation record. The low weight for antivirulence was due to the small number of samples, which was explained by the poor test coverage [32–34].

**Fig. 3.**
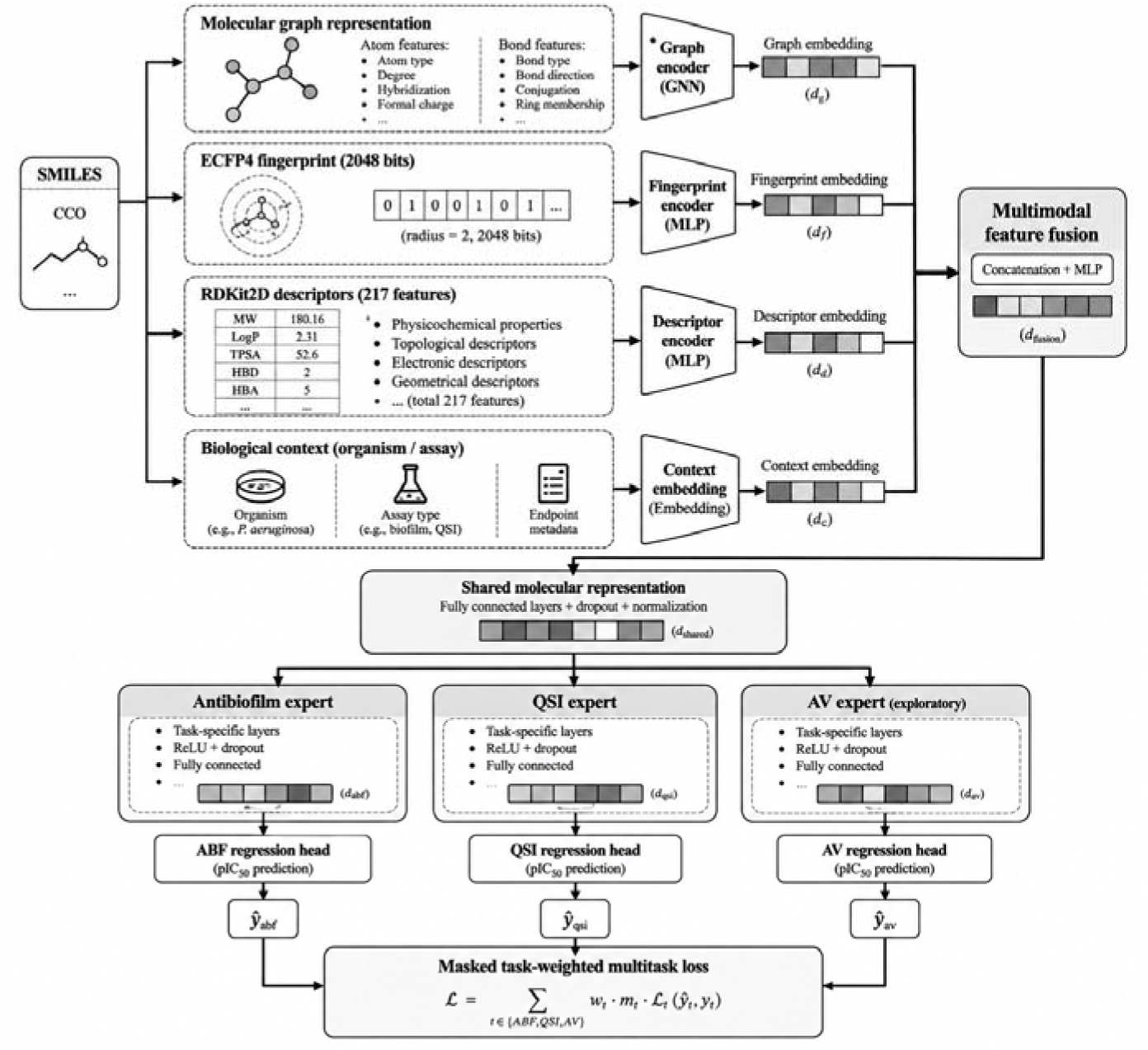
Functional schematic of graph, fingerprint, descriptor, and context fusion with task-specific regression experts.

The implemented method included molecular graph inputs, fingerprints, descriptors and biological context, and 28-dimensional atom features and task-weighted regression heads. The validated summary of the final run is documented, but the training configuration (graph depth, embedding sizes, optimizer settings, scheduling, batch size, early-stopping rule, random seeds, and software versions) are not documented in the summary. These unreported parameters are not derived from the schematic and precise checkpoint-level replication relies on the original training configuration and execution logs.

The aim can be formulated as L = Σ_t w_t Σ_(i∈O_t) Huber(y_it, ŷ_it) / |O_t| for task t and observed labels i of the set O_t, with w_t indicating the task weight and Huber(y_it, ŷ_it) representing the Huber loss function. In this formulation, task labels that are missing are not interpreted as having a negative or zero value. The actual implementation and checkpoints are the authoritative description of dimensions and optimizer configuration.

**Algorithm 3.** Masked multimodal task-expert training

1. Construct molecular graphs, descriptors, fingerprints, and context from the frozen split.
2. Fit normalizers and categorical vocabularies using training records only.
3. Encode each modality and fuse learned representations.
4. Route fused representations through task-specific experts.
5. Compute masked Huber losses with the documented task weights.
6. Optimize training loss and select architecture by primary validation score.

### 2.8. Endpoint-adaptive frozen prediction policy

A candidate task-expert model and a gated mixture-of-experts (GatedMoE) alternative were compared using the primary validation RMSE over the two adequately represented endpoints. The frozen evaluation validation score favoured TaskExpert-MTL (0.6450) over GatedMoE-MTL (0.6605). Endpoint-specific blending then combined the selected neural predictor with same-split classical models using validation-only weight search. The frozen policy used 50% neural and 50% RF for antibiofilm; 100% neural for QSI; and 30% neural plus 70% RF for the exploratory antivirulence task. Fig. 4 and Algorithm 4 describe this separation of model selection from independent testing.

**Fig. 4.**
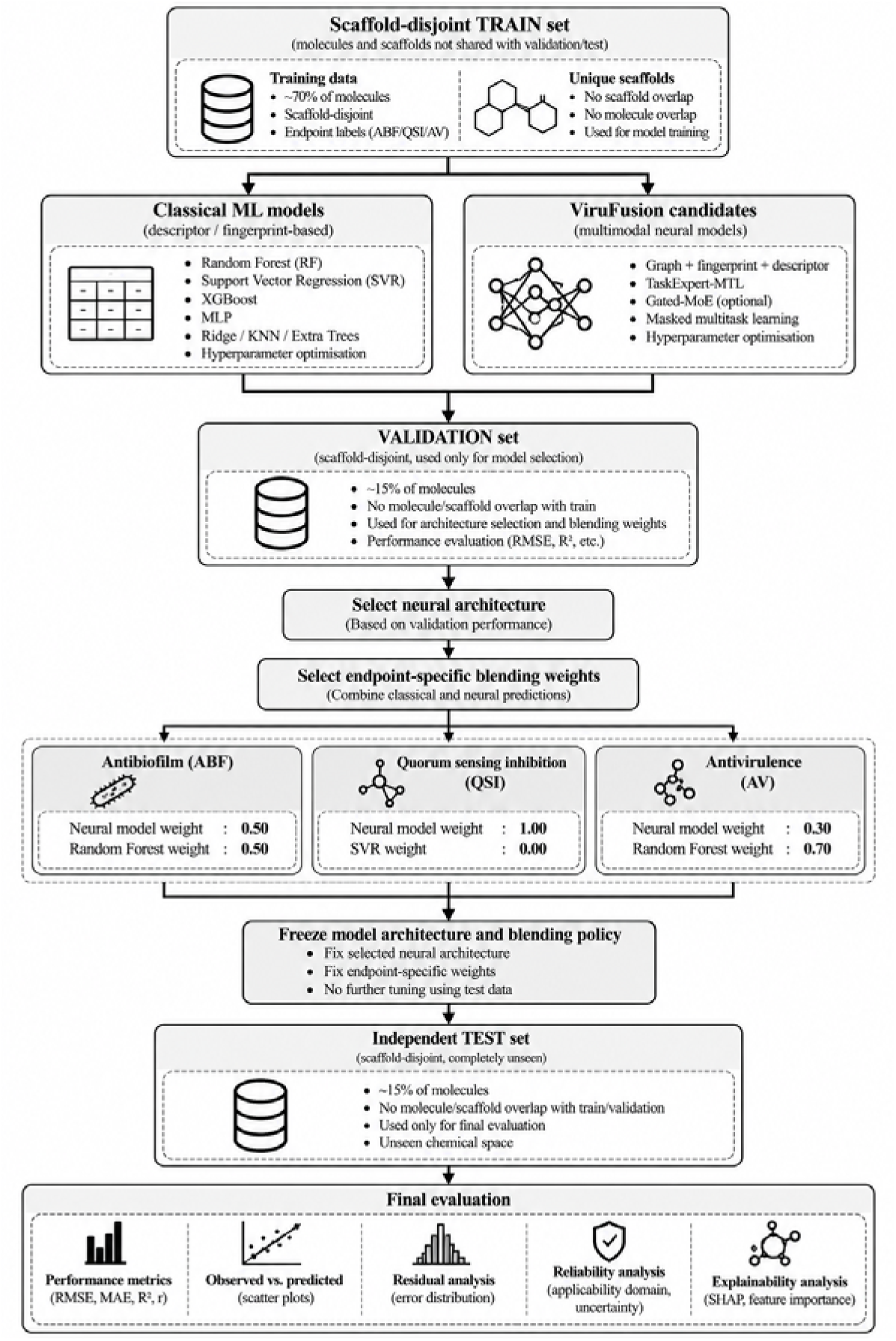
Endpoint-specific policy selection and final evaluation sequence; all selection precedes test evaluation.

The validation-selected neural contribution was fixed separately for each endpoint before independent-test evaluation: α_ABF = 0.50, α_QSI = 1.00, and α_AV = 0.30. Consequently, the QSI policy relied entirely on the neural model, while antibiofilm and exploratory antivirulence predictions were combined with their respective classical regressors. This comparison evaluated endpoint-specific fusion weights; it did not isolate the contribution of every individual representation or expert component.

For each endpoint t, the deployed policy is ŷ_t = α_t ŷ_neural,t + (1 − α_t) ŷ_classical,t, with α_t selected using validation predictions. No architecture, hyperparameter, or α_t was selected on independent-test labels.

**Algorithm 4.** Endpoint-adaptive selection and model freezing

1. Select the neural architecture using the pre-specified primary validation score.
2. Refit approved classical baselines using the same training partition.
3. Evaluate candidate convex mixture weights on validation observations.
4. Select a weight separately for each endpoint and freeze the full policy.
5. Evaluate the independent test once and report every endpoint, including failures.

### 2.9. Evaluation, statistical assessment, and reporting boundaries

Regression performance was assessed using RMSE, MAE, R^2^, Pearson correlation, Spearman correlation, and concordance correlation coefficient, as supported by the evaluation outputs. The final results reported here focus on the metrics documented in the frozen independent-evaluation summary. Grouped and paired bootstrap procedures formed part of the pipeline, but numerical confidence limits for the frozen final model were not contained in the available summary. Therefore, no p-values or inferential confidence intervals are reported. Development-stage uncertainty diagnostics were examined separately, and molecule-level observed-versus-predicted plots were not reconstructed without the corresponding frozen prediction vectors.

Statistical comparison between predictors was framed in terms of paired prediction errors on the same held-out observations. The grouped-bootstrap framework accounts for dependence among repeated observations by resampling molecular groups and recalculating paired RMSE differences. Because the numerical paired-bootstrap outputs and frozen model prediction vectors were not available in the evaluated results summary, the reported improvements remain descriptive; statistical superiority was not inferred from the point estimates alone.

The antivirulence test contains two observations. Correlation coefficients computed from two points are unstable and provide no meaningful evidence of predictive generalization; this endpoint is therefore discussed descriptively. Applicability-domain, conformal-interval, and attribution analyses are represented in the earlier development workbook but are not automatically transferable to the final frozen policy. They are treated as developmental diagnostics rather than as independently verified final optimization findings.

## 3. RESULTS

### 3.1. Curated collection and quantitative eligibility

The curated master contains 8,765 evidence records: 5,848 antibiofilm, 2,615 QSI, and 302 antivirulence observations. Only 2,959 observations satisfied the final quantitative modelling criteria. Fig. 5 shows the disparity between evidence availability and model-ready measurements. Antibiofilm contributes 2,345 model-ready records, QSI 569, and antivirulence 45. Thus, biological evidence volume cannot be equated with quantitative sample size, and the sparse antivirulence task remains a fundamental limitation. Table 2 gives the precise endpoint-level reduction, while Table 3 reports the resulting modelling imbalance.

**Fig. 5.**
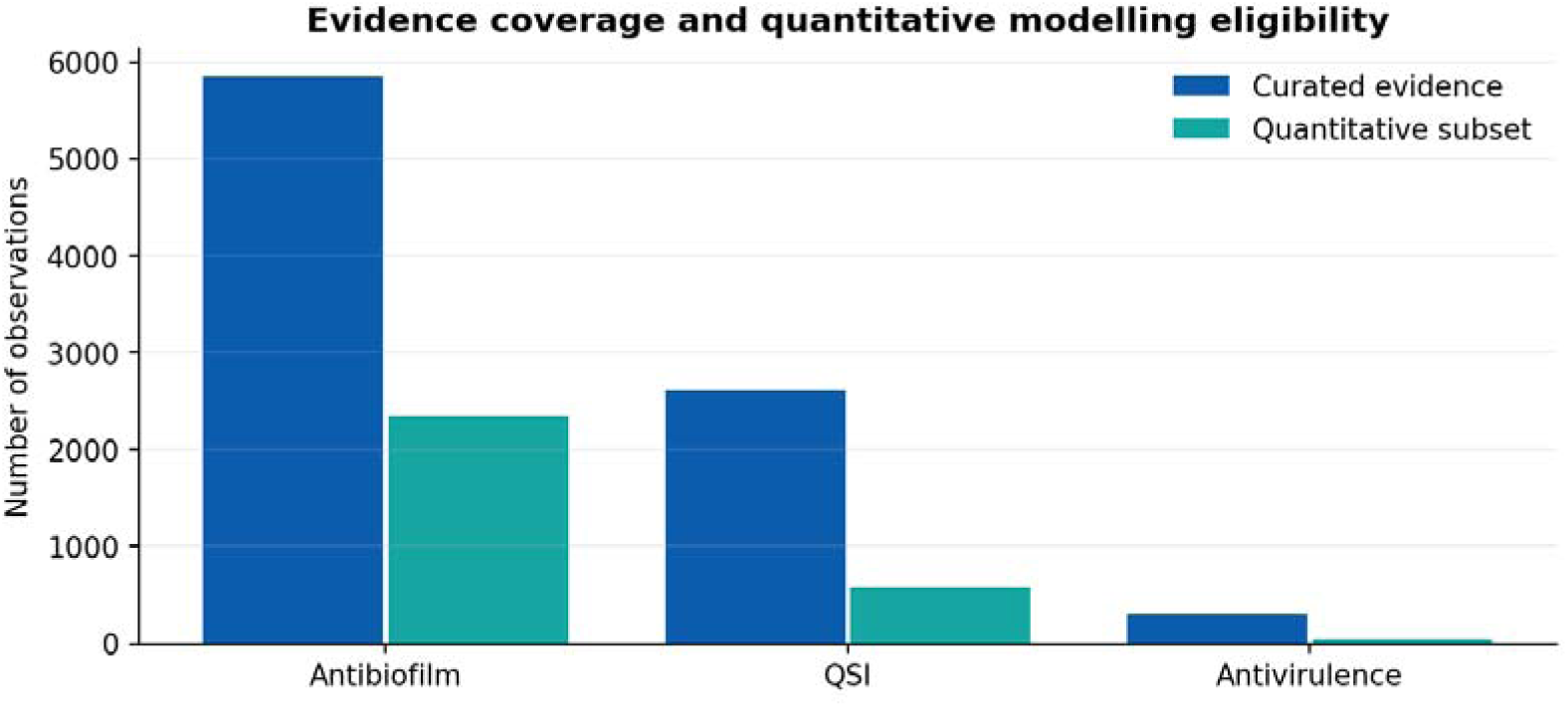
Observed curated versus model-ready endpoint counts; counts from the curated CSV and full final-run log.

### 3.2. Molecular physicochemical distribution

The selected evidence molecules contain molecules having different molecular weight, lipophilicity, and polar surface area. The plots for molecular weight, Log P, and topological polar surface area have been done in histogram format in Fig. 6. All these values have been calculated from the curated data set containing 8,765 records. The extreme points for all the distributions have been excluded from the plot only for visual clarity; however, these extreme points have been included in the actual data.

**Fig. 6.**
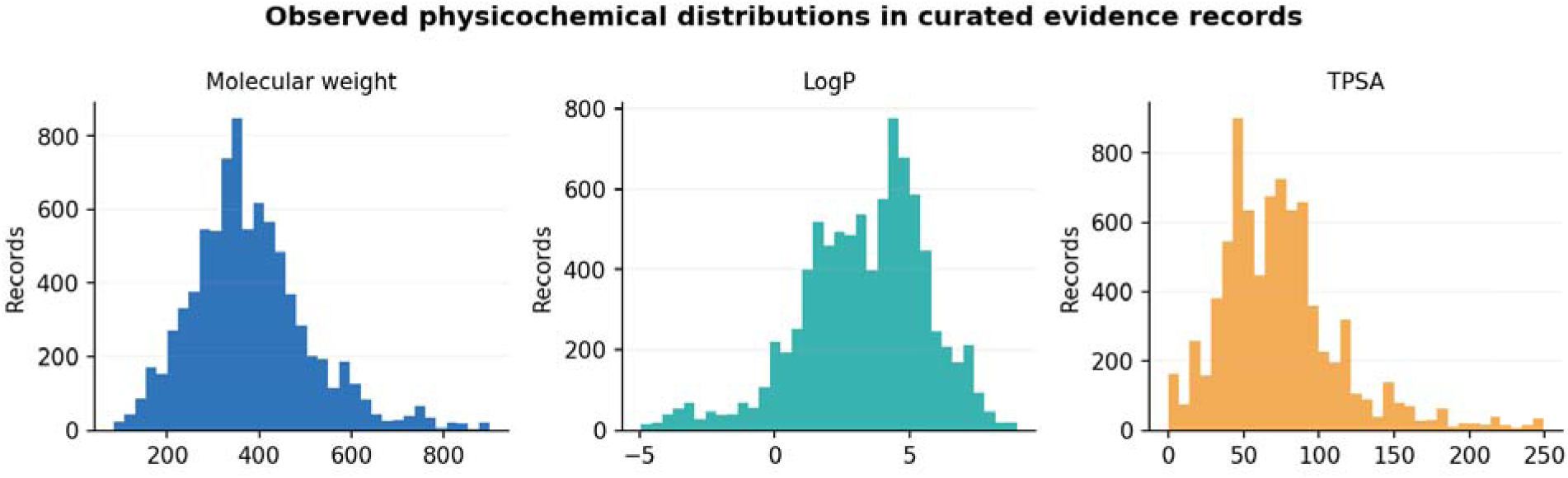
Evidence-weighted molecular-weight, LogP, and TPSA histograms from the curated master; plotting ranges are restricted for readability.

### 3.3. Scaffold composition and pathogen coverage

Multiple scaffold family types have been observed in the curated data set, thus validating the importance of scaffold-based partitioning. Figure 7 describes the top six scaffold strings of non-empty Bemis–Murcko scaffolds in the evidence records. The label represents the rank and not the actual chemical structure; scaffold information is still provided in the curated CSV file. Another imbalance can be seen in the pathogen distribution (Figure 14), where *Staphylococcus aureus* and *Pseudomonas aeruginosa* are among the biggest pathogen classes, along with records labeled as Unknowns.

**Fig. 7.**
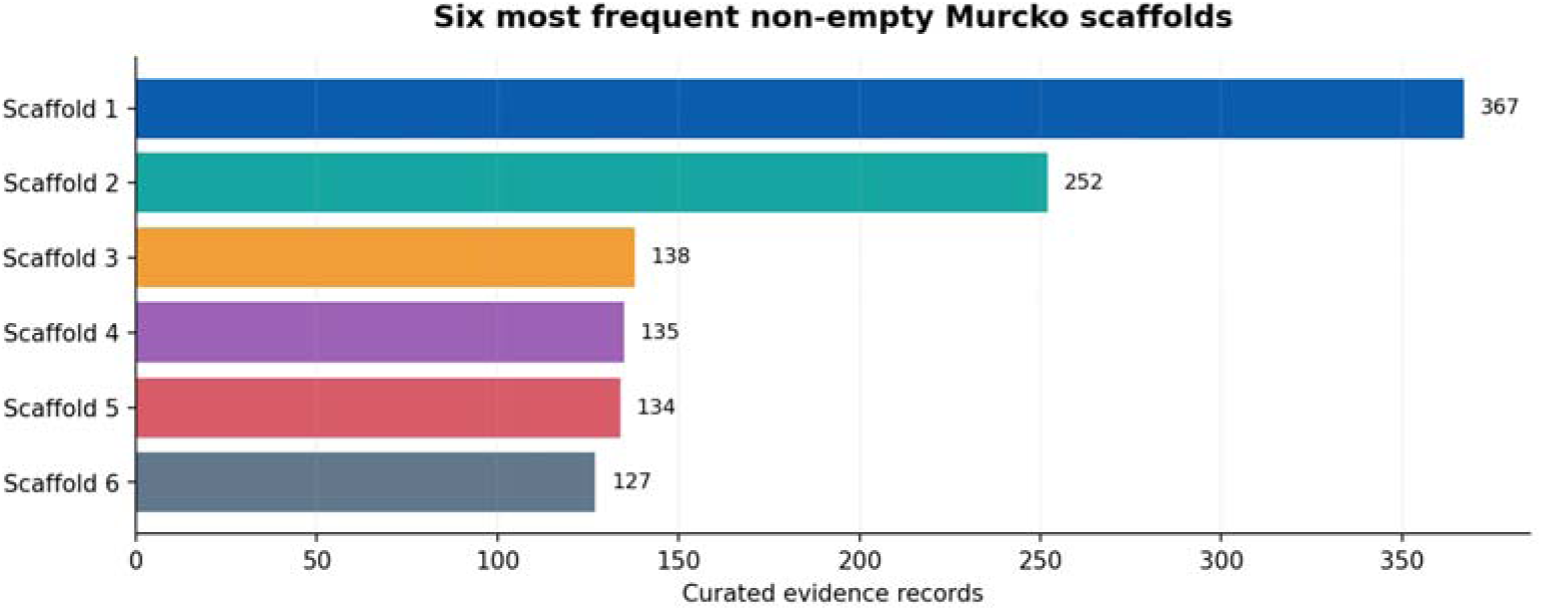
Observed frequencies of the six most common non-empty Murcko scaffolds; ranks refer to structures recorded in the curated CSV.

### 3.4. Frozen partition and leakage audit

The primary partition assigned 1,871/188/286 antibiofilm observations, 444/58/67 QSI observations and 24/19/2 antivirulence observations to Train/Validation/Test, respectively (Table 5). The resulting sample imbalance is shown in Fig. 8. The frozen evaluation audit showed that for all partition pairs there was no molecule overlap or scaffold overlap. This is a confirmation of the integrity of the primary split, but not external prospective generalization.

**Fig. 8.**
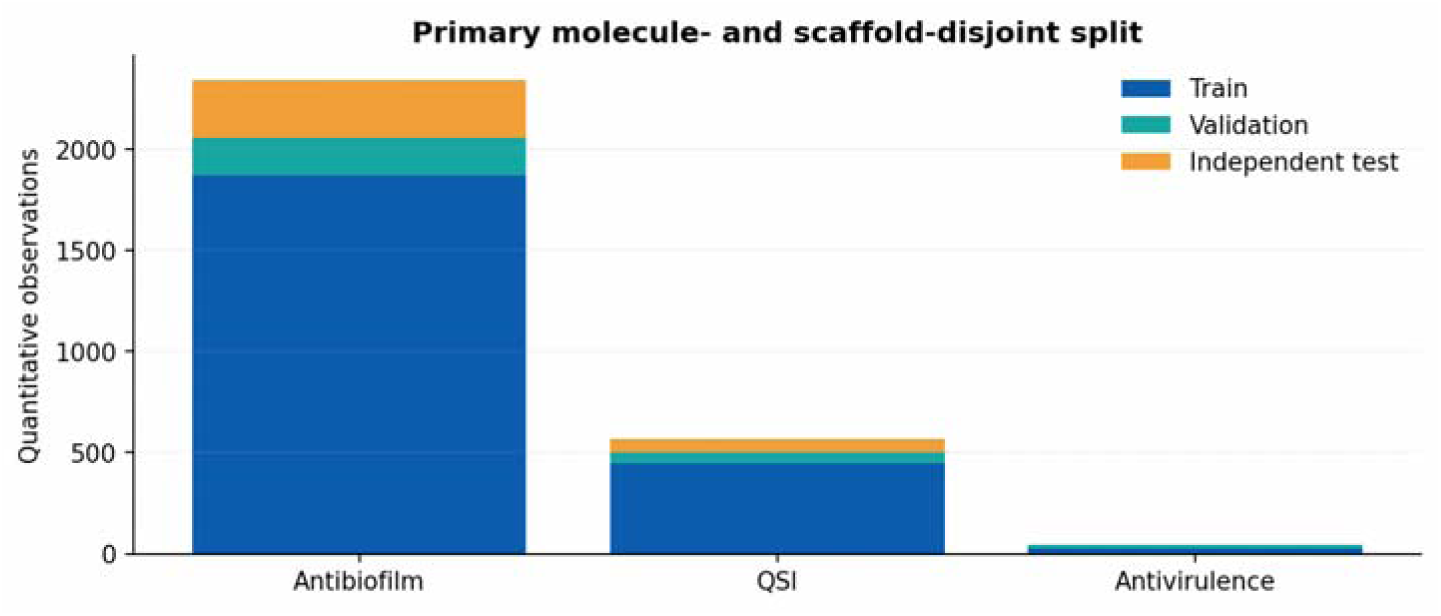
Train/validation/independent-test counts for each endpoint in the primary scaffold-disjoint split.

### 3.5. Classical QSAR performance

The benchmark classical models for the final runs were Random Forest for antibiofilm and SVR for QSI, both based on ECFP4 fingerprints. The RMSE values of their same split independent test sets were 0.6373 and 0.5794, respectively. Fig. 9 shows the endpoint-specific benchmark models. Since endpoints differ in their label distributions and test set sizes, the RMSE values were interpreted endpoint-by-endpoint and not as direct comparisons of phenotypic task hardness. Table 7 provides the same-split comparisons found in the frozen evaluation summary, but not the tuning grids of candidate models.

**Fig. 9.**
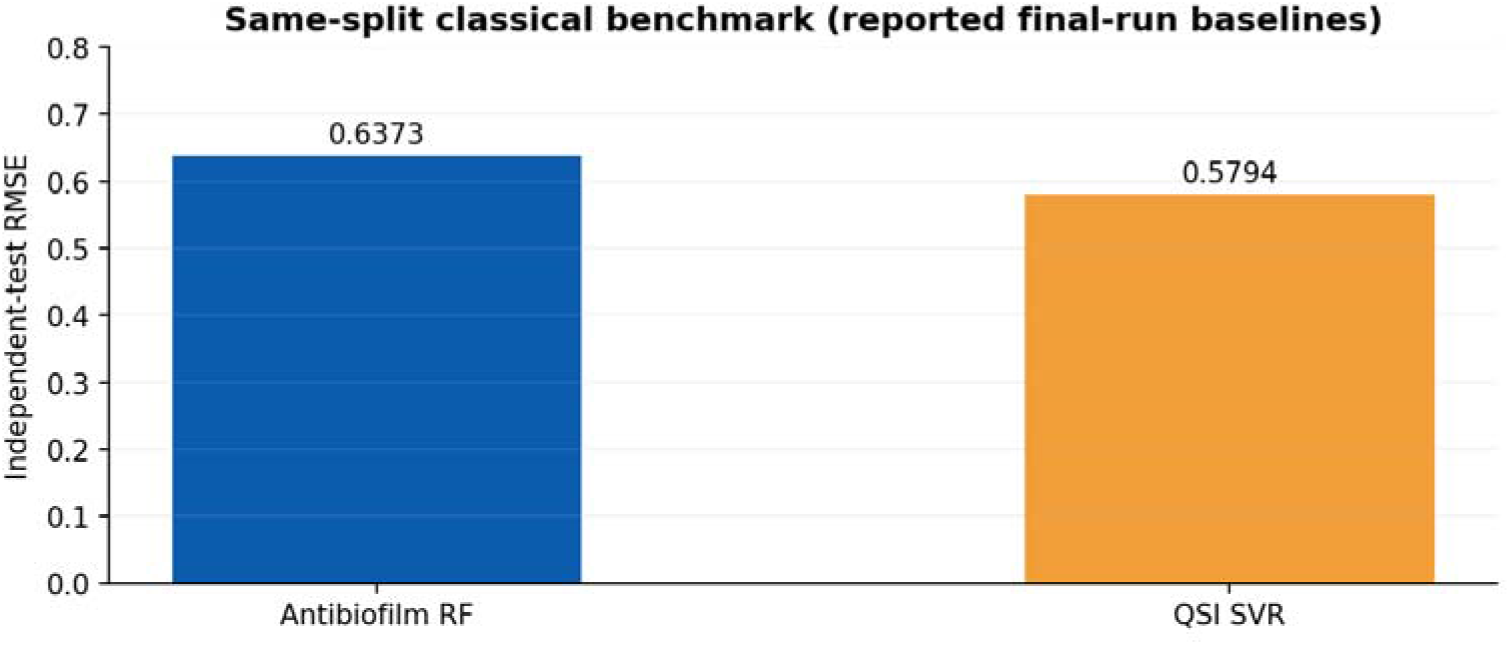
Same-split classical RMSE reference values for the two primary prediction tasks.

The antibiofilm benchmark RMSE value of 0.6373 and the QSI benchmark RMSE value of 0.5794 refer to different endpoint-specific prediction tasks. As such, the comparison was made between the respective benchmark model and the endpoint-specific final predictor only on their respective held-out test sets. Aggregate RMSE differences do not imply organism-, scaffold family-, or assay subtype-level superiority.

### 3.6. Neural architecture selection and negative transfer

The validation-only primary scores were 0.6450 for ViruFusion-TaskExpert-MTL and 0.6605 for ViruFusion-GatedMoE-MTL. The lower value was sufficient justification to select the task-expert model prior to the final independent-test evaluation. Fig. 10 provides the two validation values. The earlier ViruFusion shared development run had a QSI test RMSE of 0.6439, while the final endpoint-adaptive TaskExpert policy had a RMSE of 0.5744; Fig. 12 shows this descriptive progression. Because the figures come from separate runs, the comparison is not a controlled within-run component ablation.

**Fig. 10.**
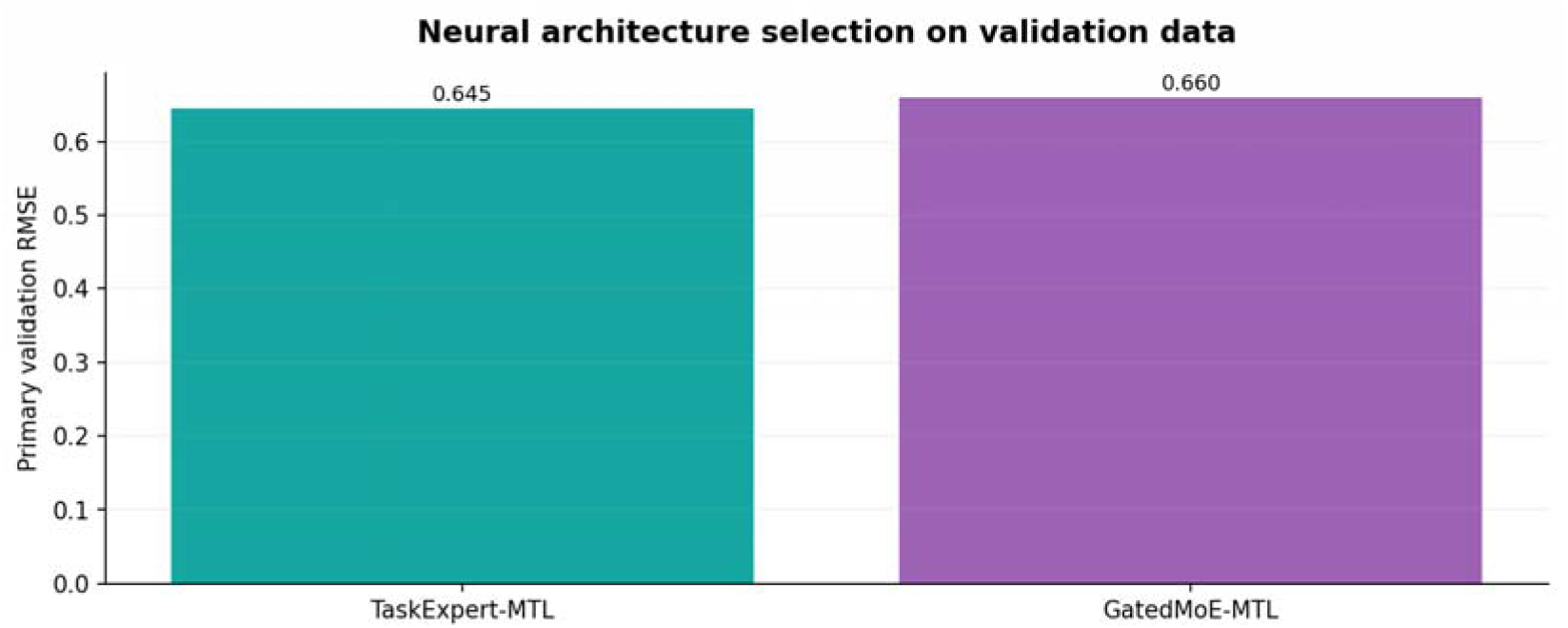
Validation-only primary RMSE comparison of the two final-run neural candidates.

### 3.7. Frozen independent-test evaluation

The independent test was tested, once architecture and blending weights were set. The endpoint-adaptive system yielded RMSE 0.5460, R^2^ 0.4816, and Pearson r 0.7101 for the antibiofilm activity (*n*=286). For QSI (*n*=67), the corresponding values were 0.5744, 0.4082, and 0.6871. These outcomes and the exploratory result on antivirulence are presented in Table 9. The two most important endpoints are shown in Fig. 11, with the metrics reported as aggregates. The value of RMSE 1.7726 and R^2^ −0.3972 produced by Antivirulence regression on two test observations did not provide generalizable prediction.

**Fig. 11.**
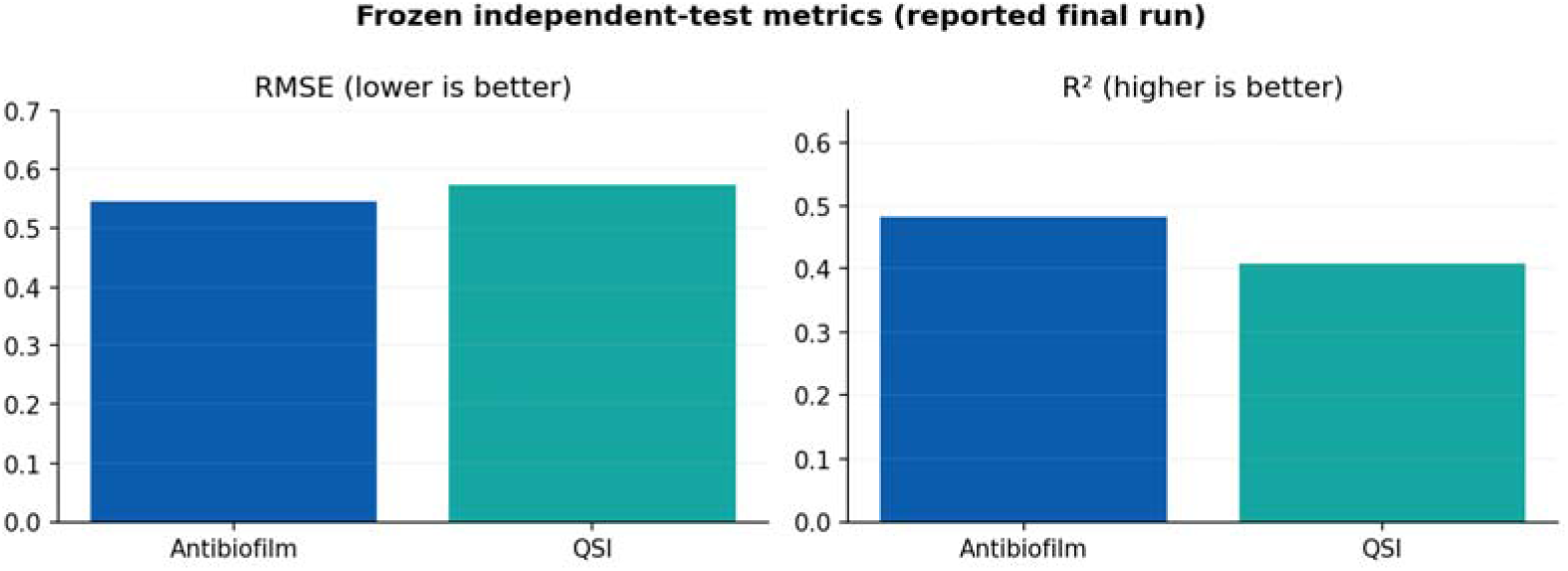
Frozen independent-test RMSE and R^2^ of the two primary tasks; aggregate metrics rather than reconstructed molecule-level predictions.

**Fig. 12.**
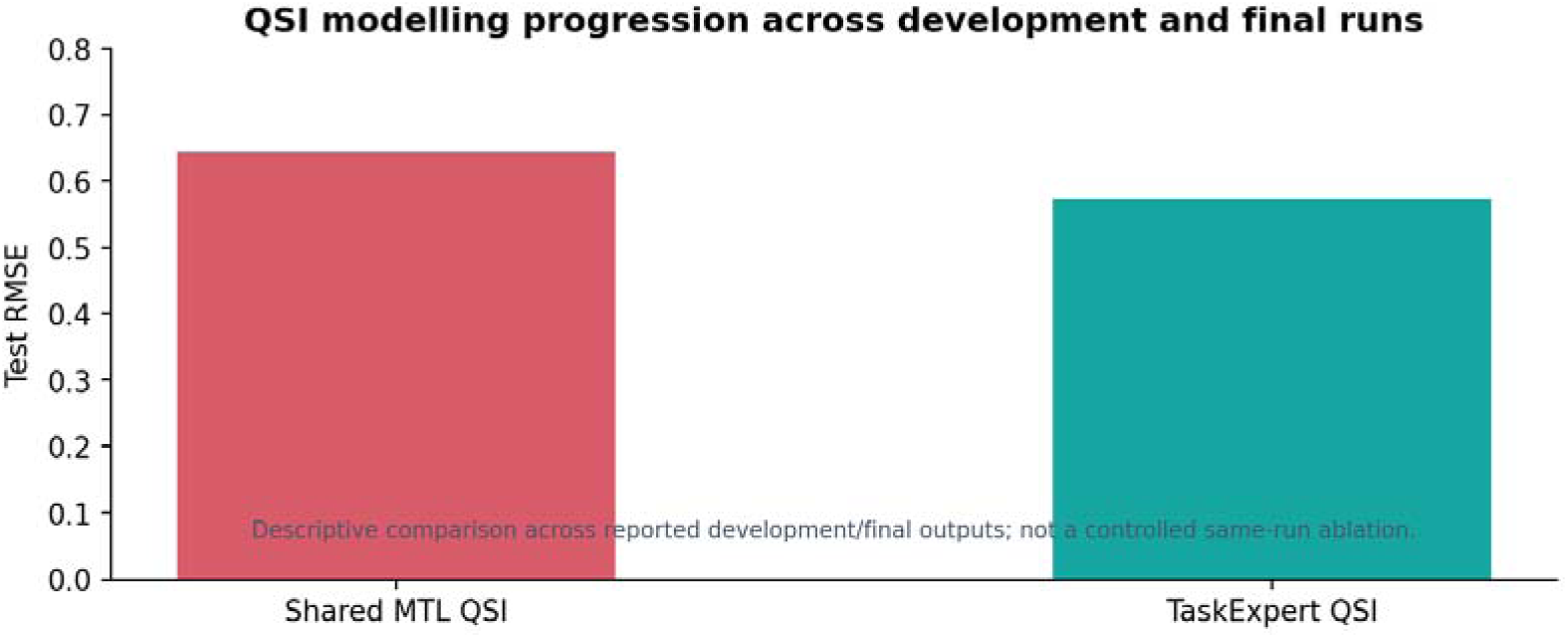
QSI modelling progression across separate development and final runs; not a within-experiment component ablation.

**TABLE 9.** Frozen independent-test regression outcomes.

| **Endpoint** | **Test *n*** | **Final policy** | **RMSE** | **R<sup>2</sup>** | **Pearson r** |
|---|---|---|---|---|---|
| Antibiofilm | 286 | 0.50 neural + 0.50 RF | 0.5460 | 0.4816 | 0.7101 |
| QSI | 67 | Neural only | 0.5744 | 0.4082 | 0.6871 |
| Antivirulence | 2 | 0.30 neural + 0.70 RF | 1.7726 | −0.3972 | Not interpretable |

***Note****: MAE, Spearman correlation, concordance correlation coefficient and bootstrap confidence limits are not tabulated because the corresponding numerical values were not present in the frozen evaluation summary used for this report.*

The RMSE reductions expressed as a percentage of the RMSE reference values were computed as 100 × (RMSE_reference − RMSE_final)/RMSE_reference. This resulted in the approximate reduction of 14.3% in antibiofilm and 0.9% in QSI. The improvement in antibiofilm was greater than the reduction in QSI was relative to the reference error. Both reductions are estimates of the value that describe a specific reduction without a paired confidence interval or an external prospective test. The antivirulence result (*n*=2) was kept only as exploratory model output due to poor test coverage, as it was not generalizable enough to assess generalizable performance.

The final system decreased the antibiofilm error by around 14.3% (0.6373 to 0.5460) compared to the reported same-split classical RMSE. For QSI, the reduction was approximately 0.9% (0.5794 to 0.5744). The difference was statistically insignificant without a matched paired confidence interval, and it was very small numerically. The RMSE for endpoints are compared in Fig. 13.

**Fig. 13.**
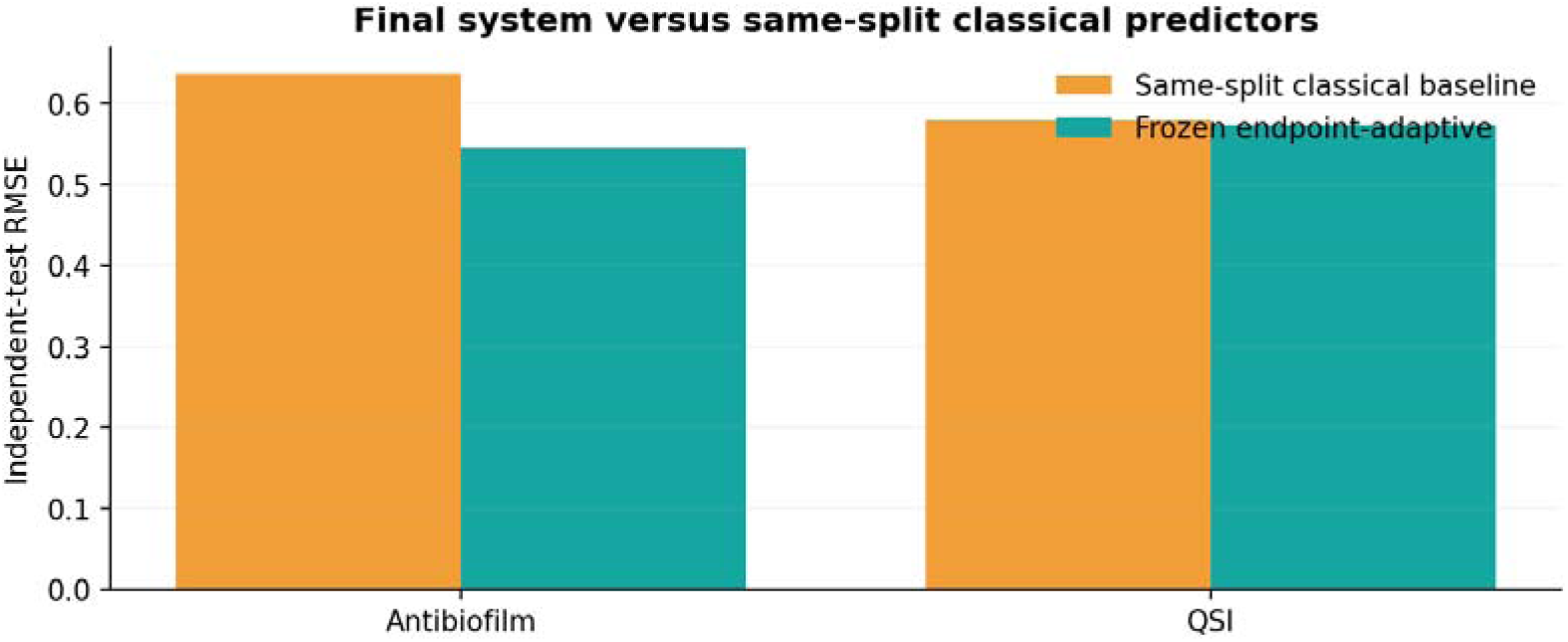
Independent-test RMSE comparison between the frozen endpoint-adaptive system and the reported same-split classical models.

**Fig. 14.**
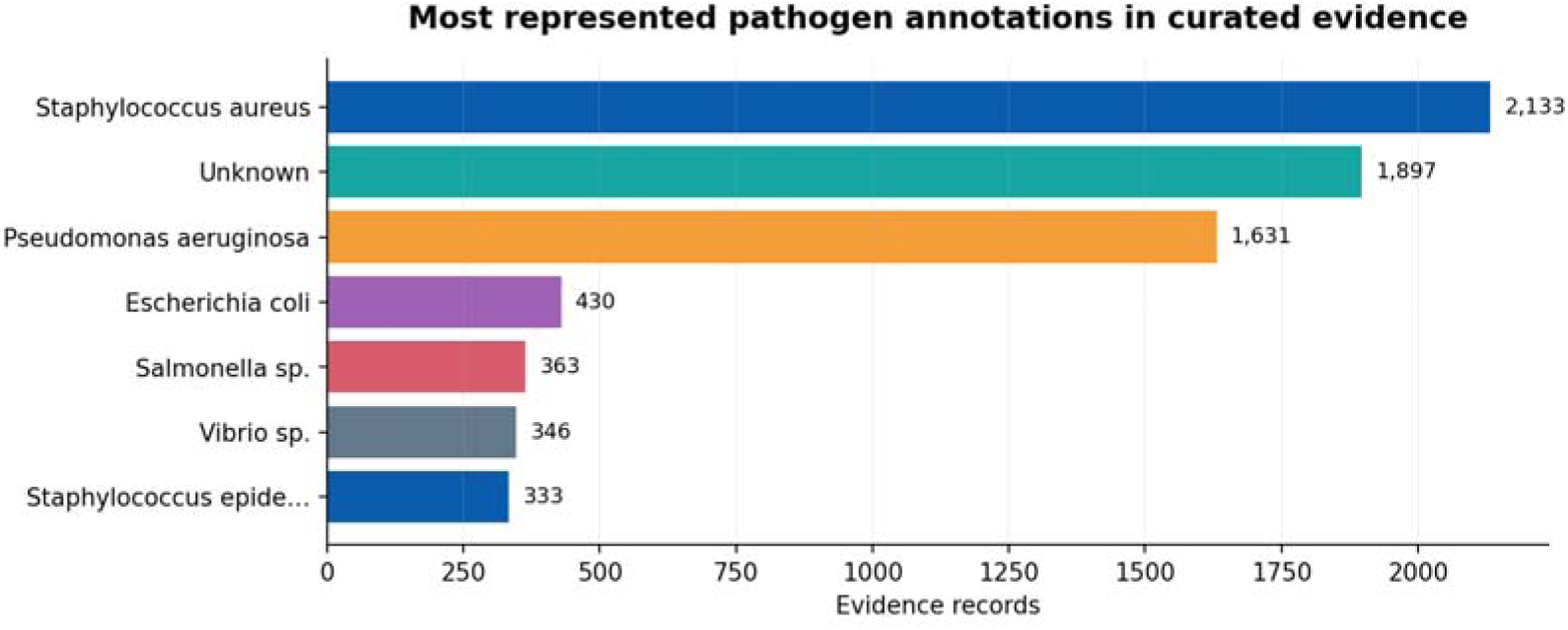
Seven most frequent pathogen annotations in the curated evidence collection, including records with unknown organism labels.

### 3.8. Representation and model-development comparisons

The previous development workbook assessed the ECFP-MLP and shared multimodal ViruFusion under the main splitting, with applicability-domain, conformal-interval and descriptor-attribution diagnostics. These were not treated as controlled ablations of the final frozen model, but rather provided for the computational development. Table 10 distinguishes between descriptions of the architecture selection, which are performed directly, and descriptions of architecture across runs. There was no improvement reported that could only be achieved if the graph or the description of the graph was altered, and no improvement reported that could only be achieved if the context was altered. Fig. 12 does not depict individual component contributions; rather it depicts progression from development to final.

**TABLE 10.** Architectural and ablation evidence status.

| Comparison | Observed evidence | Interpretive boundary |
|---|---|---|
| TaskExpert vs GatedMoE | Validation 0.6450 vs 0.6605 | Direct final-run selection |
| Shared MTL vs final TaskExpert (QSI) | RMSE 0.6439 vs 0.5744 | Across runs; descriptive only |
| Fingerprint MLP vs shared fusion | multimodal model development experiments | Component-specific effect not estimated |
| Graph-only / descriptor-only / no context | Not established in the reported evaluation | Descriptive comparison; paired CI not reported |
| Blend vs individual predictors | Final ABF/QSI RMSE plus baseline | Paired CI needed for significance |

The reported architecture comparison has been done between TaskExpert and GatedMoE under the primary validation criterion (Table 8). No matched feature-removal analysis of ECFP4-only, RDKit-descriptor-only, graph-only, no-organism-context, shared-head and complete TaskExpert variants was reported. Hence, the observed difference in validation does not negatively impact the chosen architecture at the system level but does not imply distinct numerical effects for graph encoding, global descriptors, context from the biological domain or task-specific experts.

### 3.9. Statistical reliability and interpretability

Bootstrapping tests grouped and in pairs have been included in the evaluation procedure. Confidence intervals numerically cannot be reported since they cannot be validated against the frozen final results. Thus, Table 11 differentiates documented analyses and quantitative validations in particular for the moderate improvement in QSI.

**TABLE 11.** Reliability and statistical evidence available for the frozen model.

| **Assessment** | **Verified status** | **Interpretation** |
|---|---|---|
| Leakage audit | Zero molecule/scaffold conflicts | Confirmed |
| Final grouped bootstrap | Generated in frozen evaluation record; numbers not available for quantitative reporting | CI not reported |
| Paired bootstrap | Generated in frozen evaluation record; numbers not available for quantitative reporting | No significance claim |
| Applicability domain | Available in earlier development workbook | Not final frozen-model result |
| Conformal prediction | Available in earlier development workbook | Not final frozen-model coverage |
| Explainability | Development attribution data exist | No attribution result is claimed for the frozen model |

The applicability-domain analyses on the development stage checked the structural proximity to training compounds through the ECFP4 Tanimoto similarity measure. The smaller similarity score corresponds to the less structural support for the extrapolation; however, no held-out-error-calibrated similarity threshold was found for the final frozen predictor. Conformal interval analyses have been restricted to the development stage results; the coverage could not be transferred to the final endpoint-adaptive predictor. These analyses show diagnostic potential but not calibrated uncertainty intervals for the final predictor.

Other development analyses also included Tanimoto similarity applicability domain check, conformal intervals, and descriptors attribution. They can be useful for the model development, but the coverage and the attribution depend on the model and cannot be transferred to the final frozen predictor. Figure 13 contains only test comparison results; Fig. 14 contains only independently reproducible biological annotations distribution.

### 3.10. Biological annotations, molecular interpretation, and resource utility

The evidence resource kept pathogen names, as well as assay context, allowing evaluation of biological coverage and its limitations. The seven most common pathogen labels for the curated master are shown in Fig. 14. Substantial Unknown category indicates incomplete annotation of organisms and restricts pathogen-specific interpretation. Descriptor/substructure attribution was considered to be “model sensitivity” instead of experimental evidence of biochemical target engagement.

In this study, the curated molecular evidence resource and computational prediction framework ViruDB were assessed. The reported validation did not include a publicly released, searchable interface. The experimental activities and model-predicted potencies were considered as two sets of information and were interpreted separately from the data. The evaluated software scope excludes features like compound search, interactive filtering of assays, retrieval of provenance and applicability-domain display.

Table 12 gives directly calculated molecular-property summaries of the curated evidence. There are assets for documented data and model resources as well as functions visible to the public, but not addressed in this work, which are distinguished in Table 13. As such, construction of the database and computational evaluation are presented as separate from any reference to deployment or operation to the public for search.

**TABLE 12.**
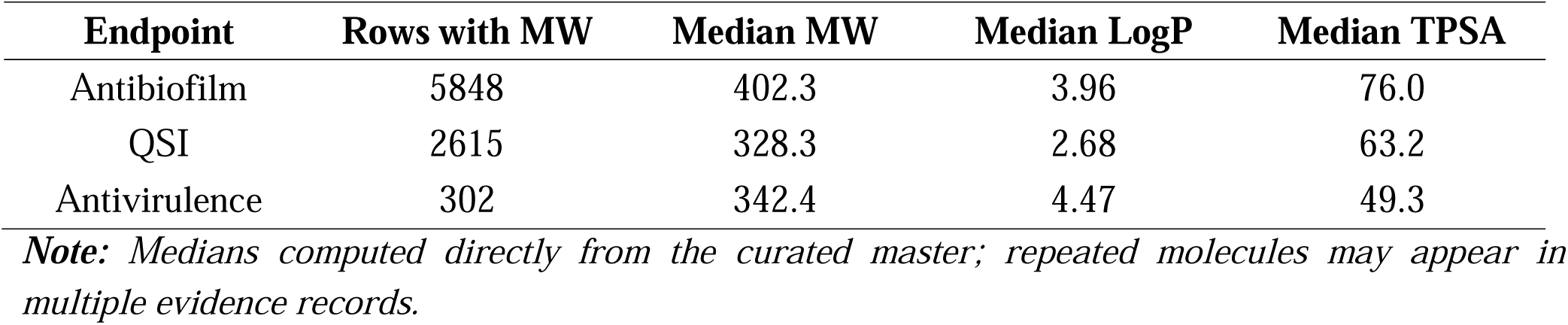
Observed evidence-weighted physicochemical summaries.

**TABLE 13.**
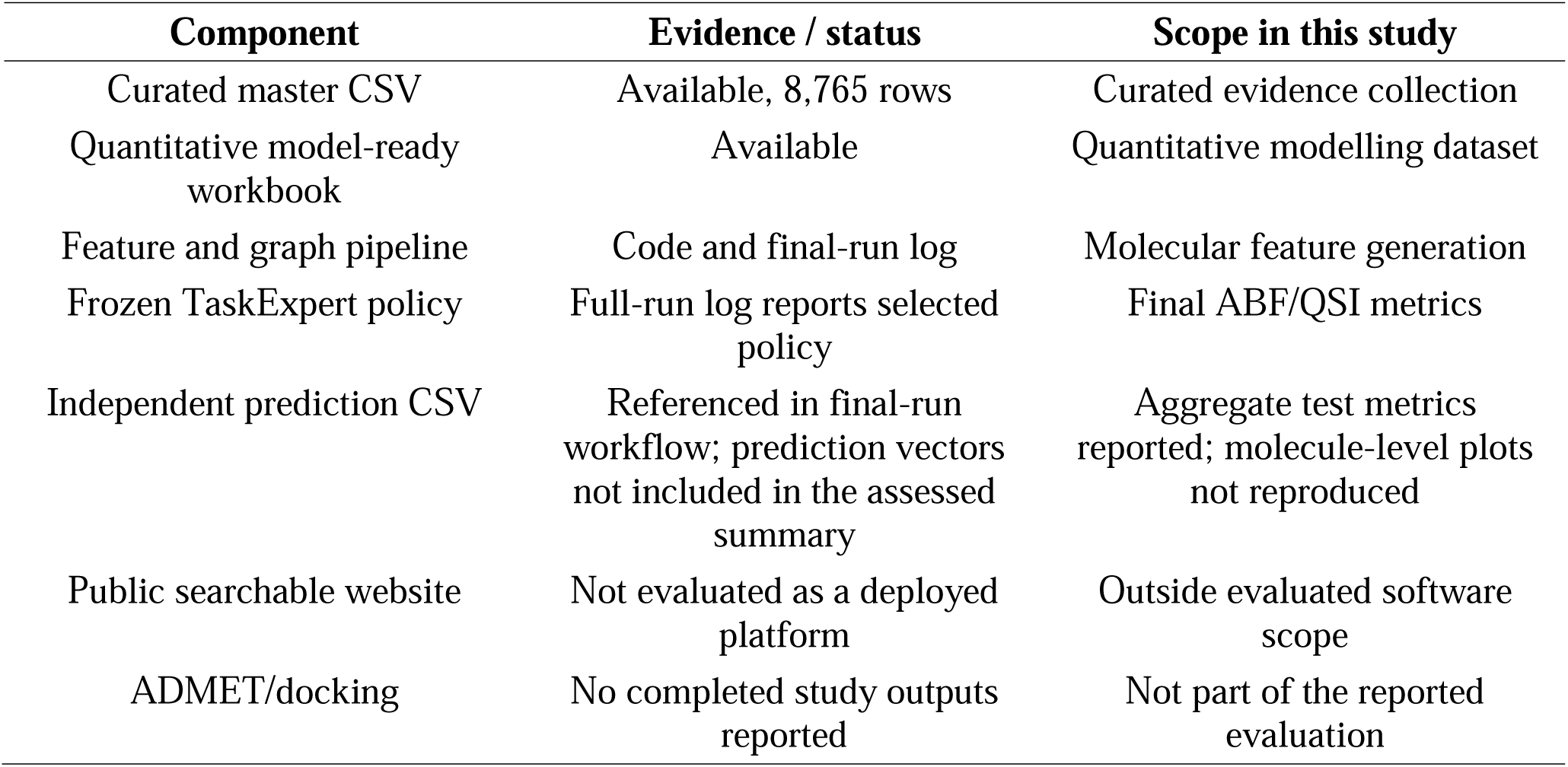
Current ViruDB research assets and implementation boundaries.

| **Component** | **Evidence / status** | **Scope in this study** |
|---|---|---|
| Curated master CSV | Available, 8,765 rows | Curated evidence collection |
| Quantitative model-ready workbook | Available | Quantitative modelling dataset |
| Feature and graph pipeline | Code and final-run log | Molecular feature generation |
| Frozen TaskExpert policy | Full-run log reports selected policy | Final ABF/QSI metrics |
| Independent prediction CSV | Referenced in final-run workflow; prediction vectors not included in the assessed summary | Aggregate test metrics reported; molecule-level plots not reproduced |
| Public searchable website | Not evaluated as a deployed platform | Outside evaluated software scope |
| ADMET/docking | No completed study outputs reported | Not part of the reported evaluation |

## 4. DISCUSSION

### 4.1. Evidence harmonization as the primary data contribution

The ViruDB results provide an example of the power of anti-virulence informatics beyond simply compiling chemical structures. The curated collection has significantly more observations than the quantitative modelling subset, in part because of the essential nature of assay eligibility and interpretable potency. The inclusion of all four of these features, biofilm biomass reduction, the inhibition of signaling reporters, bactericidal MIC and unrelated virulence phenotypes in a single continuous label could make it difficult to determine biological validity and may increase the sample size. Although organisms and mechanisms are not fully annotated, the curated fields facilitate future assay-specific analysis.

### 4.2. Placing relative to the existing resources

ViruDB is in addition to existing resources. The anti-virulence compounds, combined with pathogen and virulence factor knowledge, are incorporated into VFDB 2025, while datasets related to biofilms, prediction, and biofilm related to Anti-Biofilm, Molib, Biofilm-i, and Anti-Biofilm are emphasized. This is an assay-aware multi-endpoint evidence, a structure-based representation learning, and a documented molecule/scaffold-disjoint evaluation. Endpoint definition, species composition, and split design differences prevent one being able to draw conclusions on direct superiority over published systems from cross-paper RMSE or correlation. There would need to be a future shared benchmark in order to make a comparison of “head-to-head.

### 4.3. Negative transfer and endpoint specific learning

The validation-selected TaskExpert architecture performed better than the GatedMoE candidate on the pre-specified primary score. This discovery enables the use of task-specific representations for endpoints with the same chemical inputs but different biological measurements. This across-run comparison is descriptive, not causal, but the final QSI RMSE was less than with the previous shared multitask development model. The endpoint-adaptive selection varied between tasks as well: For the anti-biofilm task, the neural–RF mixture was selected, while the neural model was used for the QSI task. Such findings suggest that an optimal fusion weight is not consistently observed for all the heterogeneous anti-virulence-related outcomes.

### 4.4. Magnitude of predictive improvements, and limits thereof

The antibiofilm test RMSE gave an improvement relative to the reported same-split RF reference, but no matched confidence interval was provided in the frozen evaluation summary. The absolute improvement in performance for QSI was 0.0050 RMSE units, which is very limited evidence of a substantive performance advantage. For both tasks, the R^2^ is less than 0.5, meaning there is a significant amount of unexplained variability in the data even though there are positive relationships between predictions and observations. The reported performance facilitates computation of compound ranking and prioritization as an aid to, but not a substitute for, experimental screening.

### 4.5. The generalization, reliability and few antivirulence labels

Overlap of molecules and scaffolds is minimized, which adds to the strength of the evaluation. However, an internally tested non-scaffolded test is not valid for the future in other laboratories, assay technologies or pathogens. The data is highly imbalanced, and there are many cases with unreported pathogen labels. Most of all, the antivirulence endpoint is only represented by two independent test observations, thus, the regression statistics observed here are descriptive. The anti-virulence evidence collection is scientifically valuable as a curated resource, but is not sufficient for independent model validation, which will need further quantitatively comparable observations.

The prospective experimental validation is another step to come after the retrospective benchmark reported here. In such a study, the compounds used would also be different from those used in the model fit, the concentrations used would be different than those used in a subsequent assay, the bacterial-growth controls would be different, biological replicates would be used, and orthogonal confirmatory readouts would be used, excluding compounds that are not chemically related. These experimentally determined activities would be compared to those previously frozen predictions to evaluate the transfer to independent biological settings and to differentiate the inhibition of general growth from the inhibition of antivirulence activities.

### 4.6. Use an algorithm to explain and interpret biological data

Descriptor and atom-level explanations can provide ideas about the factors that might be involved in lipophilicity, polarity, charge distribution, and scaffold patterns, but offer predictive explanations, not biochemical causation. A QS reporter phenotype per se does not give any indication of binding to any specific regulator; docking scores would also be computational hypotheses if achieved. Target-engagement assays, growth controls, and orthogonal biological assays should be used to validate the mechanistic activity.

### 4.7. Limitations and future extensions of the project

This study has the following limitations: the original assays are heterogeneous, organism and target annotation is incomplete, sample numbers are small for regressing antivirulence activity, there is no external independent validation set available for public access that was independently deposited, the final frozen evaluation bootstrap was not available to the reader within the current manuscript bundle, and a validated public interface was not available. Proposed but not implemented: ADMET prediction and target-guided docking. The most important future steps that should be taken are: to publish a versioned, licence-compliant dataset and model; to reproduce statistical confidence intervals; to validate search and inference software; to increase the number of chemically novel candidates and compounds tested in prospective labs; and to produce a virus-inhibition dataset and model that is versioned and licence-compliant.

The final computational evaluation can be reproduced based on the curated and model-ready datasets, source and assay dictionaries, molecule/scaffold partition identifiers, processing objects, the selected TaskExpert checkpoint, the endpoint-specific weights, the independent-test predictions and references, the metric scripts, and a documented software environment. The current report outlines the corresponding research workflow, but not to make stable public deposits of this complete frozen asset set. To achieve public release and independent replication, the same files need to be archived using the right source-level license and source information.

## 5. CONCLUSION

ViruDB incorporates a carefully selected collection of 8765 anti-virulence related experimental evidence records, and a quantitative, scaffold-disjoint modelling benchmark of 2959 observations. The endpoint-adaptive ViruFusion framework is a molecular representation that was fused with task-specific learning and validation-selected classical–neural prediction. For the reported frozen independent test, the values of RMSE for the antibiofilm model and QSI model were 0.5460 and 0.5744, respectively. While these findings are useful for prioritization of compound selection as part of the assay process, they do not prove therapeutic activity or mechanistic target engagement for newly predicted compounds. There are still some limitations, such as the lack of antivirulence test and limited external validation. ViruDB maintains the molecular identity, the context of the assays, and the ability to evaluate the results in light of leakage, thus offering a reproducible ground for further expansion of the data, software deployment and potential anti-virulence discovery.

## CRediT authorship contribution statement

**Pankaj Paul:** Writing – review & editing, Writing – original draft, Visualization, Validation, Methodology, Investigation, Formal analysis, Supervision and Conceptualization. **Ruchita Santra:** Writing – review & editing, Formal analysis.

## Funding

No funding was received for conducting this analysis.

## Declaration of interest

The authors declare that they have no known competing financial interests or personal relationships that could have appeared to influence the work reported in this paper.

## Data and code availability

The curated molecular evidence, quantitative modelling data, and implementation workflow are described in Table 13. No stable public repository identifier or complete archived set of frozen model checkpoints, held-out prediction vectors, and execution-environment files was documented in the material evaluated for this manuscript. Accordingly, this article does not assert public access to these complete research assets. Any subsequent release remains subject to the licensing conditions of the originating data sources.

## Notes

### Competing Interest Statement

The authors have declared no competing interest.

